# Modeling Type 1 Diabetes progression from single-cell transcriptomic measurements in human islets

**DOI:** 10.1101/2023.07.19.549708

**Authors:** Abhijeet R. Patil, Jonathan Schug, Chengyang Liu, Deeksha Lahori, Hélène C. Descamps, the Human Pancreas Analysis Consortium, Ali Naji, Klaus H. Kaestner, Robert B. Faryabi, Golnaz Vahedi

## Abstract

Type 1 diabetes (T1D) is a chronic condition in which the insulin-producing beta cells are destroyed by immune cells. Research in the past few decades characterized the immune cells involved in disease pathogenesis and has led to the development of immunotherapies that can delay the onset of T1D by two years. Despite this progress, early detection of autoimmunity in individuals who will develop T1D remains a challenge. Here, we evaluated the potential of combining single-cell genomics and machine learning strategies as a prime approach to tackle this challenge. We used gradient-boosting-based machine learning algorithms and modeled changes in transcriptional profiles of single cells from pancreatic tissues in T1D and nondiabetic organ donors collected by the Human Pancreas Analysis Program. We assessed whether mathematical modelling could predict the likelihood of T1D development in nondiabetic autoantibody-positive organ donors. While the majority of autoantibody-positive organ donors were predicted to be nondiabetic by our model, select donors with unique gene signatures were classified with the T1D group. Remarkably, our strategy also revealed a shared gene signature in distinct T1D associated models based on different cell types including alpha cells, beta cells and acinar cells, suggesting a common effect of the disease on transcriptional outputs of these cells. Together, our strategy presents the first report on the utility of machine learning algorithms in early detection of molecular changes in T1D.

## Introduction

Muscle and adipose tissues respond to insulin to increase glucose uptake. Insulin is a hormone which is made by specialized cells called beta cells positioned in the islets of Langerhans in the pancreas. In the autoimmune disease Type 1 Diabetes (T1D), which arises from a complicated interplay between genetic and environmental factors, T cells attack and destroy beta cells. During the early stages of the autoimmune process, autoantibodies (AAb) against pancreatic islets can frequently be detected in the serum and the presence of multiple autoantibodies is a strong predictor of progression to T1D^1^. Although the discovery of insulin was a milestone in T1D research that made survival of millions of patients possible, insulin therapy does not provide complete protection against diabetes-associated complications. Recent research has revealed various immune cell types and secreted cytokines responsible for beta cell destruction^2^. These findings have led to the development of therapies to slow down or prevent the T1D onset. In particular, blocking T cells using Teplizumab was recently approved by FDA and has been reported to delay progression to clinical T1D in high-risk participants by two years^3^. Despite this breakthrough and the promises of ongoing immunotherapy trials for T1D, the unmet clinical need is to reliably identify individuals fated to develop T1D at the earliest possible stages and substantially delay or prevent the disease onset.

One promising approach to unravel early molecular events leading to beta cell destruction is to generate molecular maps of individual cells from tissues relevant to the etiology of T1D and rigorously compare them across organ donors at different disease stages. The JDRF-supported nPOD^4^ (https://npod.org/) and the NIDDK-supported HPAP consortia^5, 6^ (https://hpap.pmacs.upenn.edu) have contributed to this effort by collecting pancreatic tissues and immune-related organs from hundreds of non-diabetic, but islet autoantibody-positive (AAb+), and T1D organ donors. Among numerous genomics and molecular assays, the revolutionizing single-cell transcriptomics (scRNA-seq) has become a standard technology to study T1D development. The first set of human donor islets analyzed by the HPAP team released transcriptional profiles of islets in 24 non-diabetic (CTL), AAb+, and T1D donors across more than ∼80,000 cells^7^. In this study, we reported a surprising correlation between the expression level of around 1,000 genes in beta cells, but not any other cell types, with anti-GAD autoantibody levels detected in the serum of AAb+ donors, suggesting that the progression of the autoimmunity process is reflected in the transcriptome of AAb+ beta cells. Despite the new insights gained from scRNA-seq profiling in this study and other reports^8^, many questions related to the early molecular events leading to autoimmunity in T1D remain unanswered. For example, it is not clear if there are consensus transcriptional changes associated with T1D in different islet cell populations across the human population. In addition, it remains unknown whether there are any consensus transcriptional changes associated with T1D progression which can be detected at early stages of autoimmunity in AAb+ donors.

Here, we aimed to address these questions by modeling T1D progression using machine learning approaches on an expanded dataset of scRNA profiling from 50 HPAP organ donors. We reasoned that machine learning strategies, which can learn patterns from data, may identify consensus patterns of change in gene expression associated with T1D at prediabetic stages. A machine learning model is trained to perform a task by receiving several examples of input data, such as gene counts as features, with corresponding output labels, e.g., individual cells labelled as T1D, AAb+, or CTL. The model then updates internal parameters to enhance prediction accuracy. We devised a machine learning classifier based on the eXtreme Gradient Boosting^9^ (XGBoost) algorithm and carried out classifications of single cells across the three donor groups. Remarkably, we report that T1D can be modeled by this machine learning algorithm with high accuracy using solely islet cell transcriptomic data from T1D and CTL donors. Interestingly, our classifier reported T1D-like islet cells in a subset of AAb+ donors, demonstrating that the transcriptional adaptations that occur in islets of T1D patients are already initiated in some AAb+ donors.

## Results

### Single-cell RNA sequencing data in human pancreatic islets

We took advantage of the most recent release of scRNA-seq experiments by HPAP across 50 donors in three groups namely T1D (n=9), AAb+ (n=10), and CTL (n=31) (Fig. 1a). The preprocessing of scRNA-seq data included filtering low-quality cells, doublet removal, and dimensionality reduction, similar to our previously described protocol^10, 11^. We annotated individual cells based on the expression of known marker genes using scSorter^12^ and reported the frequency of 10 different cell types across donors (Fig. 1b-f). Overall, acinar, alpha and beta cells were the largest cell populations with 43401, 47988, and 36837 cells, respectively. The percentage of acinar and alpha cells was evenly distributed across different donor groups (Fig. 1g). Expectedly, beta cells were significantly less abundant in T1D donors compared to other donor groups, reflecting the autoimmune destruction of this cell type (Fig. 1g). Conversely, ductal cells were more abundant in tissues collected from T1D donors than the other two groups, reflecting the difficulty of isolating pure islets from these donors (Fig. 1g). The cell type annotation and composition across different groups were also consistent with previously published studies^7, 10, 11^. The expression of marker genes across major cell types such as acinar (*PRSS1*), alpha (*GCG*), beta (*INS*), delta (*SST*), ductal (*KRT19*), endothelial (*VWF*), epsilon (*GHRL*), immune (*NCF2*), PP (*PPY*), and stellates (*COL1A1*) further corroborated cell annotations across our samples (Fig. 1h). Together, we have generated high-quality transcriptional data across single cells and annotated major cell types in islets of three donor groups.

**Fig. 1.**
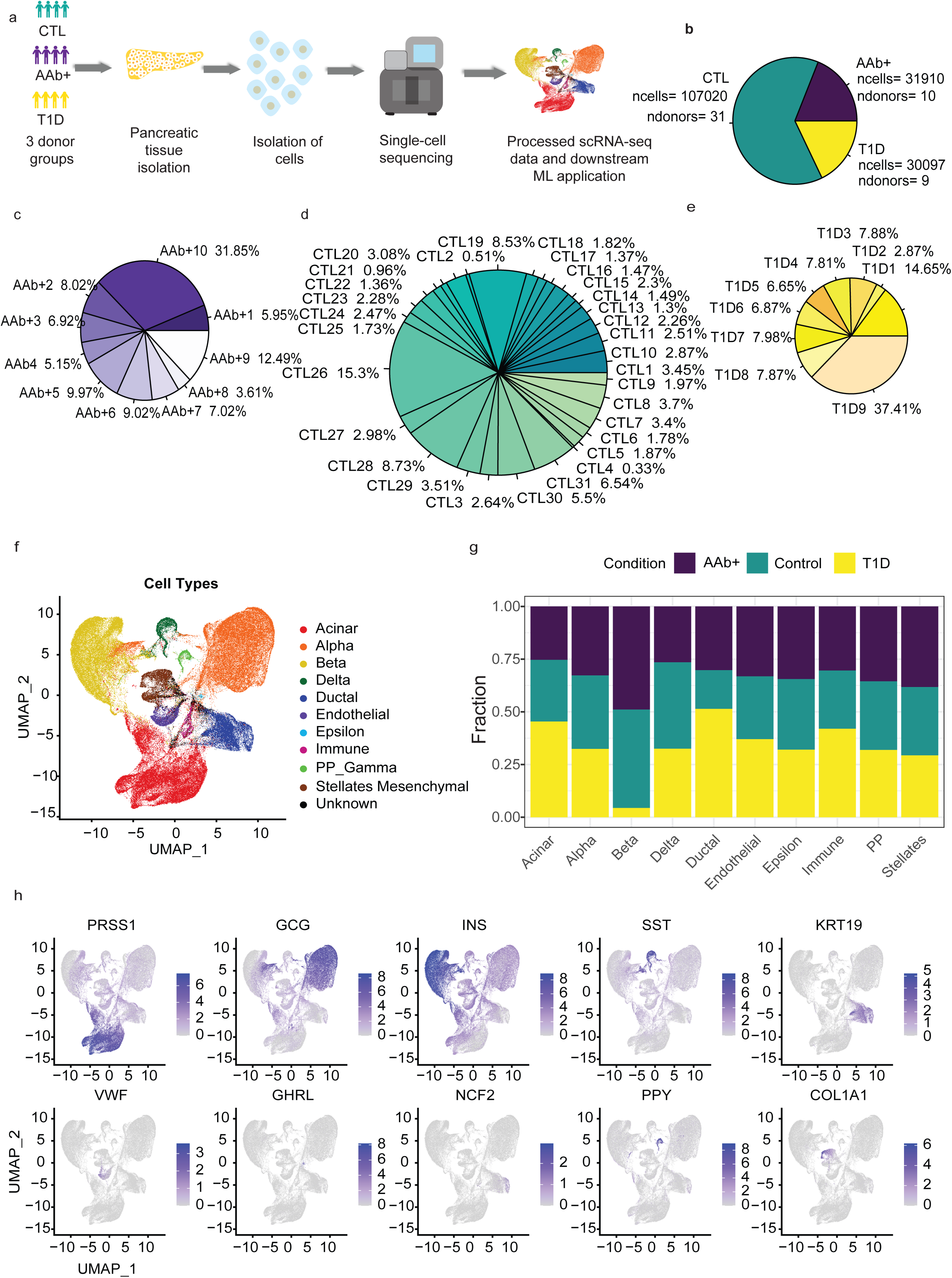
scRNA-seq reveals the cell populations of the human pancreatic islets in CTL, AAB, and T1D donors. **a.** The complete workflow depicting the scRNA-seq and machine learning workflow using human pancreatic islet tissue samples. **b.** Pie chart showing the number of cells and donor distribution across different biological conditions. **c-e.** Pie chart showing the distribution of cells across each donor. (c) AAB donors (d) Control donors (e) T1D donors. **f.** Uniform manifold approximation and projection (UMAP) plot showing the scSorter cell classification of islet cells. **g.** Stacked bar chart showing the percentage-wise distribution of cell types across AAB, Control and T1D donors. **h.** Multiple feature plots UMAPs depicting the validation of cell type-specific expression of marker genes. Acinar cells (*PRSS1* high), Alpha cells (*GCG* high), Beta cells (*INS* high), Delta cells (*SST* high), Ductal cells (*KRT19* high), Endothelial cells (VWF high), Epsilon cells (*GHRL* high), Immune (*NCF2* high), PP cells (*PPY* high), Stellates cells (*COL1A1* high).

### Performance of the machine learning model on scRNA-seq islet data

For both ‘annotated’ and ‘unannotated’ strategies, we divided individual cells into training and testing groups and subjected the training data to hyper-parameter optimization (HPO) using the XGBoost algorithm. After performing HPO using a 5-fold cross-validation procedure, we obtained the optimal parameter set, which was used for training and testing the final model. We performed 100 repetitions of the above procedure by randomly shuffling the training data (i.e., random sampling without replacement). In the ‘unannotated’ XGBoost classifier built for all cells, the T1D vs AAb+ and T1D vs CTL binary classifiers performed exceptionally well, averaging ∼99% accuracy, ∼99% sensitivity, and ∼97% specificity. The AAb+ vs CTL classifier demonstrated an accuracy of ∼96% and a specificity of ∼88%. The relatively small decrease in performance in the AAb+ vs CTL comparison likely reflects the similarity in transcriptional landscapes of single cells from AAb+ and CTL donors (Fig. 2b and Table S1).

**Fig. 2.**
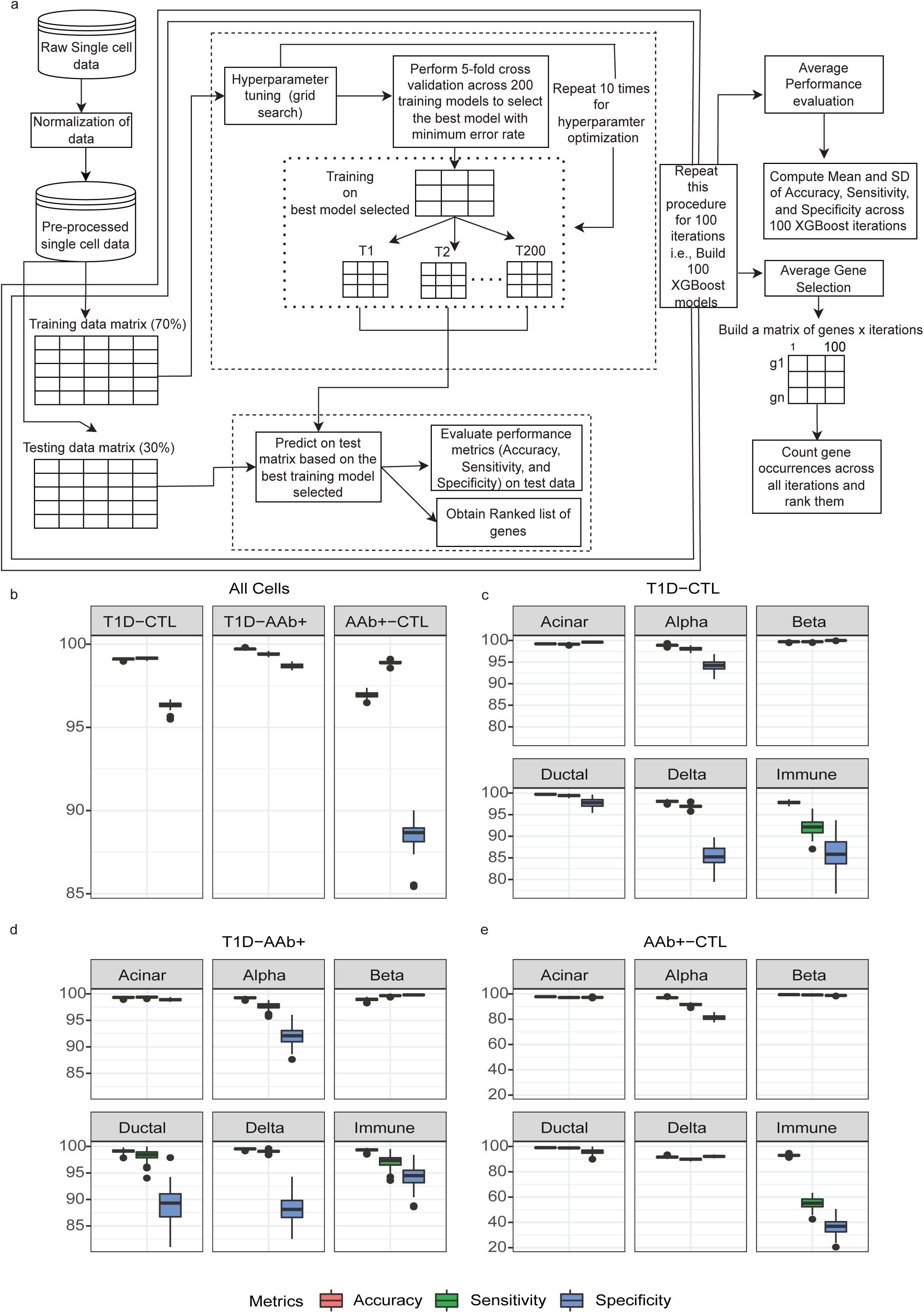
Classification performance of machine learning model on scRNA-seq islet data. **a.** A schematic workflow of the XGBoost and performance. The ML-based XGBoost model was built for gene selection and classification. The dotted lines show the training and testing procedures, where T denotes the gradient boosting tree models. The double lines show 100 repetitions of the entire workflow. **b.** Box plots depicting a pairwise comparison of the XGBoost method across all cells in the dataset. **c.** Performance of XGBoost across major cell types for T1D vs. CTL comparison using boxplots. **d.** Performance of XGBoost across major cell types for T1D vs. AAB comparison using boxplots. **e.** Performance of XGBoost across major cell types for AAB vs. CTL comparison using boxplots.

In the ‘annotated’ classification between T1D and CTL groups built for each annotated cell type, the XGBoost method performed exceptionally well on our three metrics in the acinar, alpha, beta, and ductal cells. However, binary classification using delta cells or immune cells, which contained fewer cells compared to other cell types, demonstrated a reduced performance (Fig. 2c). Similar results were observed in the T1D vs AAb+ and AAb+ vs CTL comparisons (Fig. 2d-e). Altogether, the ‘annotated’ XGBoost classifier exhibited high performance across all cell-type comparisons (Fig. 2c-e). Additionally, the average standard deviation (SD) in the comparisons of T1D vs CTL, T1D vs AAb+, and AAb+ vs CTL across 100 repetitions were found to be very low (<1%), demonstrating the robustness and stability of XGBoost models (Table S1).

### Top-ranked genes selected from the machine learning model and pathway enrichment analysis

A major reason for our choice of XGBoost over other machine learning algorithms such as convolutional neural networks is the interpretability and transparency in the XGBoost’s decision-making process. In particular, XGBoost produces feature importance rankings, allowing us to understand which features, i.e. genes, drove predictions. In contrast, neural networks are often considered ‘black boxes’, making it challenging to interpret their predictions^13^. To better understand which features drove the high-performance predictions across single cells, we obtained the key gene signatures for each comparison and used two strategies: (1) we ranked the lists of genes based on the Robust Ranking Algorithm (RRA) approach^14^ (Table S2-S4) and (2) we examined the unranked list of genes based on their selection frequency across 100 repetitions (Table S5-S7). These top-selected genes were used for downstream pathway or protein-protein interaction analysis to further understand the difference between the three clinical conditions considered.

The ranked list of genes with P-value < 0.05 based on the RRA approach obtained from the unannotated T1D-CTL classifier were enriched with genes annotated as ‘lipid mRNA metabolic process’, ‘defense to external biotic’, and ‘antimicrobial’ pathways (Fig. 3a). The 20 KEGG pathways (FDR < 0.05) enriched within top features (ranked genes with P-value < 0.05) of unannotated and annotated T1D-CTL classifiers were related to HIV and HPV infections in addition to cytokine signaling (Figs. 3b, S1-S2). The enrichment of shared pathways across features from classifiers built based on distinct cell types such as alpha cells and ductal cells motivated us to carefully evaluate commonality among top features (i.e., genes) across different cell types. We applied the RRA approach again but now on the ranked list of genes across cell types. This strategy can produce reliable and consistent rankings even in the presence of noisy or incomplete data. In T1D vs CTL classification, while more than one hundred genes were uniquely detected in classifiers built on each annotated cell type such as acinar, alpha, ductal, and beta cells, around 66 of these genes were common across classifiers of all annotated cell types based on their high RRA scores, suggesting changes in the transcriptional outputs across T1D islets independent of their cellular ontogeny (Fig. 3c). In particular ‘neutrophil degranulation’, ‘ER-phagosome pathway’, and ‘pancreatic secretion’ were enriched within this set of 66 commons genes associated with models based on annotated cell types (Fig. 3d). Moreover, in AAb+ vs CTL and T1D vs AAb+ classifiers, both common and unique genes across different annotated cell types were detected, suggesting the relevance of multiple pathways to changes in cells from AAb+ donors (Fig. 3e-f). Together, the XGBoost classifiers based on training the model using single cells grouped as different cell types revealed the link between multiple genes and pathways associated with T1D and autoantibody positivity.

**Fig. 3.**
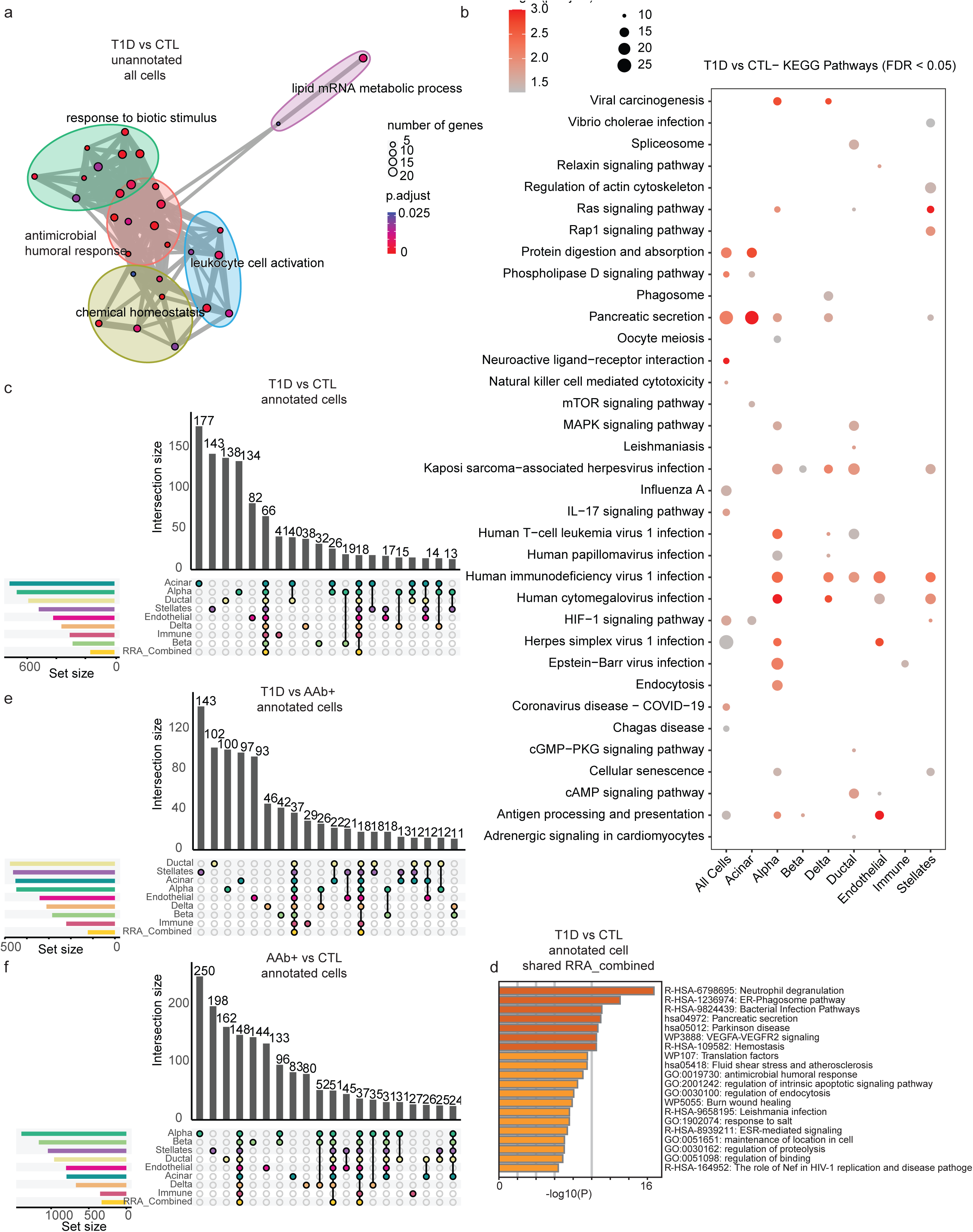
Top-ranked genes selected from the machine learning model and pathway enrichment analysis. **a.** GO cluster analysis showing pathways based on genes obtained from T1D vs. CTL comparison across unannotated all cells. **b.** Top 20 KEGG pathways based on ranked genes obtained from T1D vs. CTL comparison across Unannotated all cells and annotated different cell types (Table S2). **c.** UpSetR chart showing the common and unique count of genes across annotated cells in T1D vs. CTL comparison. **d.** Pathways based on shared RRA_combined gene list in T1D vs. CTL annotated cells. **e.** UpSetR chart showing the common and unique count of genes across annotated cells in T1D vs. AAb+ comparison. **f.** UpSetR chart showing the common and unique count of genes across annotated cells in AAb+ vs. CTL comparison.

### Expression of HLA class I genes in beta cells across healthy and T1D donors

We next focused on the beta cell classifiers within all three groups and focused on genes with a selection frequency higher than 50% following the unranked gene selection approach. In particular, we examined the enrichment of highly frequent genes as top features within KEGG pathways (Fig. 4a-b). Among the top 10 significant pathways (FDR < 0.05), the “Type 1 diabetes mellitus” and “antigen processing” were found among the highly frequent features of T1D vs CTL and T1D vs AAb+ classifiers for beta cells (Table S11-S12). The specific genes involved in Type 1 diabetes and antigen processing and presentation pathways were predominantly from HLA class I, i.e., *HLA-A*, *HLA-B*, *HLA-C*, and *HLA-E*. Additionally, the selection frequency of these genes was 100% among all 100 repetitions of modeling, suggesting that HLA class I genes were reproducible top features across donors.

**Fig. 4.**
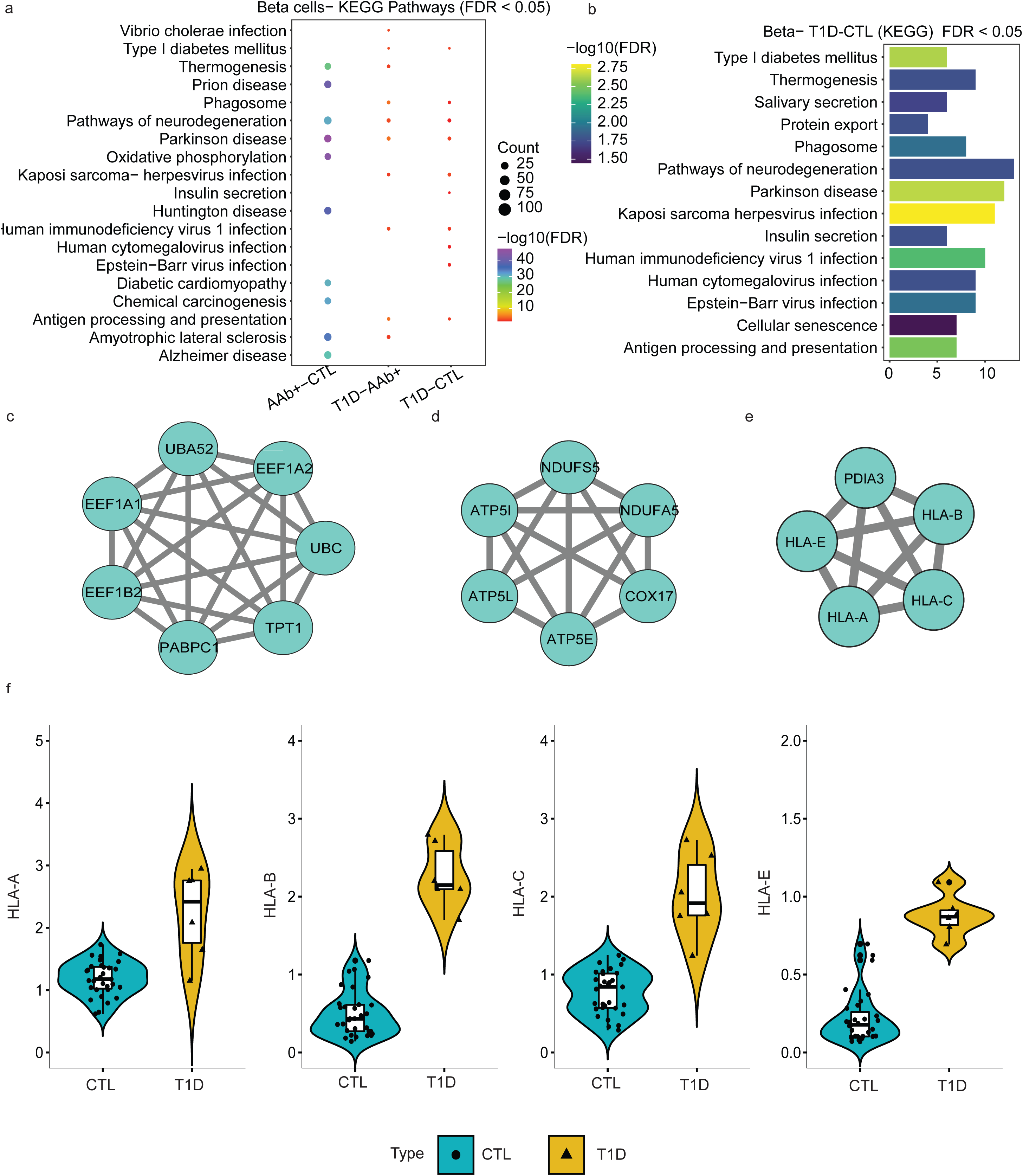
Expression of HLA-1 genes in beta cells across healthy and T1D donors. **a.** Comparison of significant KEGG pathways (FDR < 0.05) obtained from different pairwise classifiers. **b.** Significant KEGG pathways for T1D vs. CTL (FDR < 0.05). **c-e.** The top 3 modules were obtained from the PPI network using the MCODE algorithm. **a. f.** Average expression of beta cells in non-diabetic controls and type-1 diabetic donors.

Next, we created a protein-protein-interaction (PPI) network map with significant genes as nodes and their connections as edges. We further applied the MCODE algorithm^15^ on the PPI network map and obtained three clusters or key modules (Fig. 4c). All the HLA class I proteins were grouped into one cluster that was significant (P-value < 0.05). In the beta cell classification, genes encoding the HLA class I proteins were significantly upregulated in T1D donors compared to CTL donors (Fig. 4f). Another significant cluster contained genes important for mitochondrial function, while another one highlighted transcriptional elongation. Next, we analyzed the expression of the class I HLA genes in CTL and T1D beta cells individually (Fig. 4f). The dots inside the violin plots represent individual donors, where the cell-level gene counts were aggregated into pseudo-bulk counts. Strikingly, the HLA class I genes were upregulated within the few remaining beta cells from T1D donors (Fig. 4f). Together, modeling differences in beta cells of T1D and CTL donors suggest that small residues of beta cells in T1D donors express high levels of HLA class I genes. These results are in agreement with prior findings of increased HLA class I expression in islets from T1D patients obtained using antibody staining^16^.

### Prediction of AAb+ cells using classification models from annotated cells in T1D and CTL donors

One key goal in this study was to evaluate whether any subset of islet cells from AAb+ organ donors demonstrate transcriptional similarity to those from T1D individuals, and if such a similarity can be modeled, which gene features will classify a single AAb+ cell as T1D. Hence, for each annotated cell type, we used the trained T1D vs CTL classifier, where cells from T1D donors were labeled as class 1 and cells from CTL donors were labeled as class 0. We tested how this model predicted the class of AAb+ cells. Using the probability scores obtained for each individual cell from the AAb+ donor group, we determined whether the cell was predicted as T1D or CTL where a probability of > 0.5 being classified as T1D and less than 0.5 as CTL. Although 90% of cells from AAb+ donors were predicted to be non-diabetic (class 0), around 10% of cells were classified as ‘T1D’ across different cell types (class 1) (Fig. 5a). Importantly, these ‘T1D-like’ cells were not present at uniform abundance among all organ donors, but highly enriched among specific islet donors, in particular those from HPAP donors HPAP092 and HPAP107 (Fig. 5a). Strikingly, the beta cells classified as ‘T1D’ from AAb+ donors had transcriptomic signatures of HLA class I genes exactly similar to the T1D group (Fig. 5b). We also compared the expression of alpha cells from AAb+ donors classified as ‘T1D’ and observed similar results where HLA class I genes were upregulated in those AAb+ cells that were predicted as ‘T1D’ (Fig. 5c). The pancreatic alpha cells are known to have a key role in the development of diabetes mellitus^17, 18^. It has been reported that the alpha cells from donors with recent-onset T1D demonstrate reduced glucagon secretion and dysregulated gene expression^19^. Another study using immunofluorescence staining showed that the majority of HLA class I genes are expressed on pancreatic alpha cells and particularly hyper-expressed in the T1D group^20^. Further, we combined all the cells from the two AAb+ donors that were majorly predicted as ‘T1D’ and compared their HLA class I genes between all groups. We confirmed that the *HLA-A*, *HLA-B*, *HLA-C*, and *HLA-E* genes were highly upregulated in cells from AAb+ donors predicted as ‘T1D’, which might reflect that autoimmunity had already progressed in these AAb+ individuals (Fig. 5d). Of note, HPAP107 is an organ donor who was positive for both GAD and IA-2 autoantibodies, indicative of further disease progression, while HPAP092 is a single GAD autoantibody positive donor. Together, our modeling strategy discovered that a subset of cells in specific AAb+ organ donors have transcriptional patterns resembling those typically found in islets from T1D patients.

**Fig. 5.**
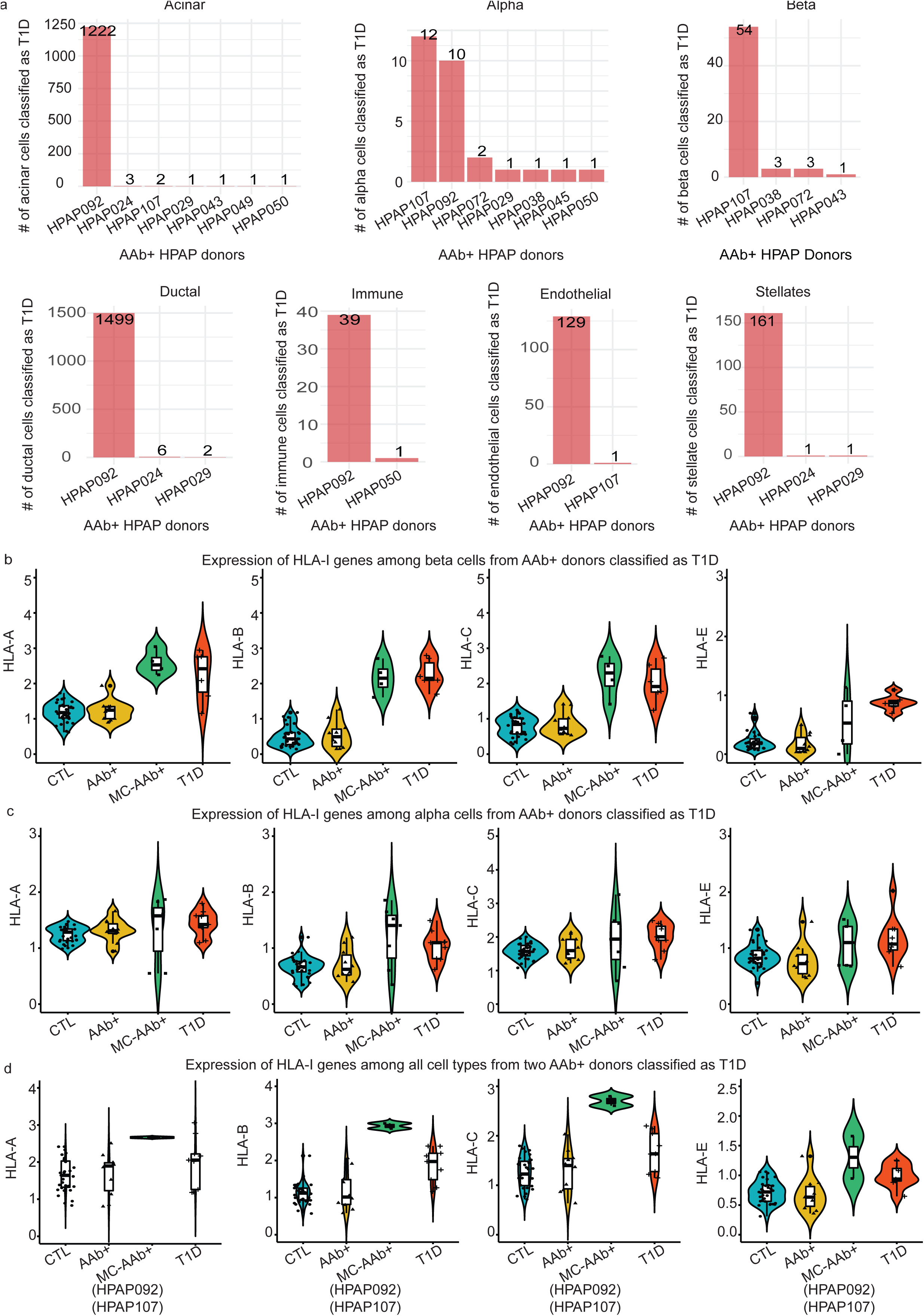
Prediction of AAB+ cells using trained T1D-CTL classifier across major cell types. **a.** Distribution of cells misclassified as T1D in different cell types. **b.** Comparing the average expression of HLA-I genes among beta cells from AAB+ donors classified as T1D with other conditions. **c.** Comparing the average expression of HLA-I genes among alpha cells from AAB+ donors classified as T1D with other conditions. **d.** Comparing the average expression of HLA-I genes among all cell types from two AAB+ donors classified as T1D with other conditions.

### Prediction of AAb+ cells using classification models from unannotated cells in T1D and CTL donors

We next compared the transcriptomic differences and similarities between T1D, AAb+, and CTL groups using our pre-trained unannotated T1D-CTL classifier. Among ten AAb+ donors, different percentages of cells from six AAb+ donors were classified as ‘T1D’, with the majority belonging to HPAP092 (Fig. 6a). We next sought to evaluate transcriptional profiles of the AAb+ cells that were predicted as ‘T1D’. To provide a reference for these ‘T1D’ AAb+ cells, we first focused on significant genes obtained in unannotated T1D vs CTL, T1D vs AAb+, and AAb+ vs CTL classifiers (Table S5-S7). We checked the gene selection frequency of HLA class I, HLA class II, and non-HLA genes across different unannotated classifiers (Fig. 6b). The selection frequency of HLA class I genes was higher compared to some of the HLA class II genes. In addition, we also checked the selection frequency of non-HLA genes including *INS, IL32, TNFAIP3,* and *LMO7* that have been associated with T1D pathology^21, 22^. The expression of HLA class I and II genes among all the AAb+ cells from HPAP092 classified as ‘T1D’ had a similar expression pattern as the T1D group (Fig. 6d-e). Previous studies have shown the inherited risk for T1D to be largely determined by specific *HLA-DQA1, HLA-DQB1, HLA-DRA, HLA-DPA1, and HLA-DRB1* alleles^21, 23, 24^. The expression of *HLA-DPB1* was found to be higher in endocrine cell types of single-cell islet data from diabetic individuals^7^. We also observed a downregulation of the *INS* gene in AAb+ cells (HPAP092), which was similar to the T1D group (Fig. 6f). Moreover, the *IL32, TNFAIP3, and LMO7* genes were found to be upregulated in AAb+ cells predicted to be in the T1D class (HPAP092). Previously, an association to T1D had been reported for these non-HLA genes (*INS*^21, 22^, *TNFAIP3*^22, 25^, *LMO7*^22, 26^, and *IL32*^27^). The similarities between the transcript levels of the HLA class I (Fig. 6d), HLA class II (Fig. 6e), and non-HLA genes (Fig. 6f) observed between AAb+ cells predicted as ‘T1D’ and the actual T1D group suggest that in specific AAb+ organ donors, the pathogenic process towards T1D had progressed further than that typical of single AAb-positive individuals. Altogether, modeling of the transcriptomic differences between islets cells from T1D and CTL donors using either an annotated or an unannotated classifier can delineate individual cells from AAb+ donors with transcriptional similarity to those from T1D patients.

**Fig. 6.**
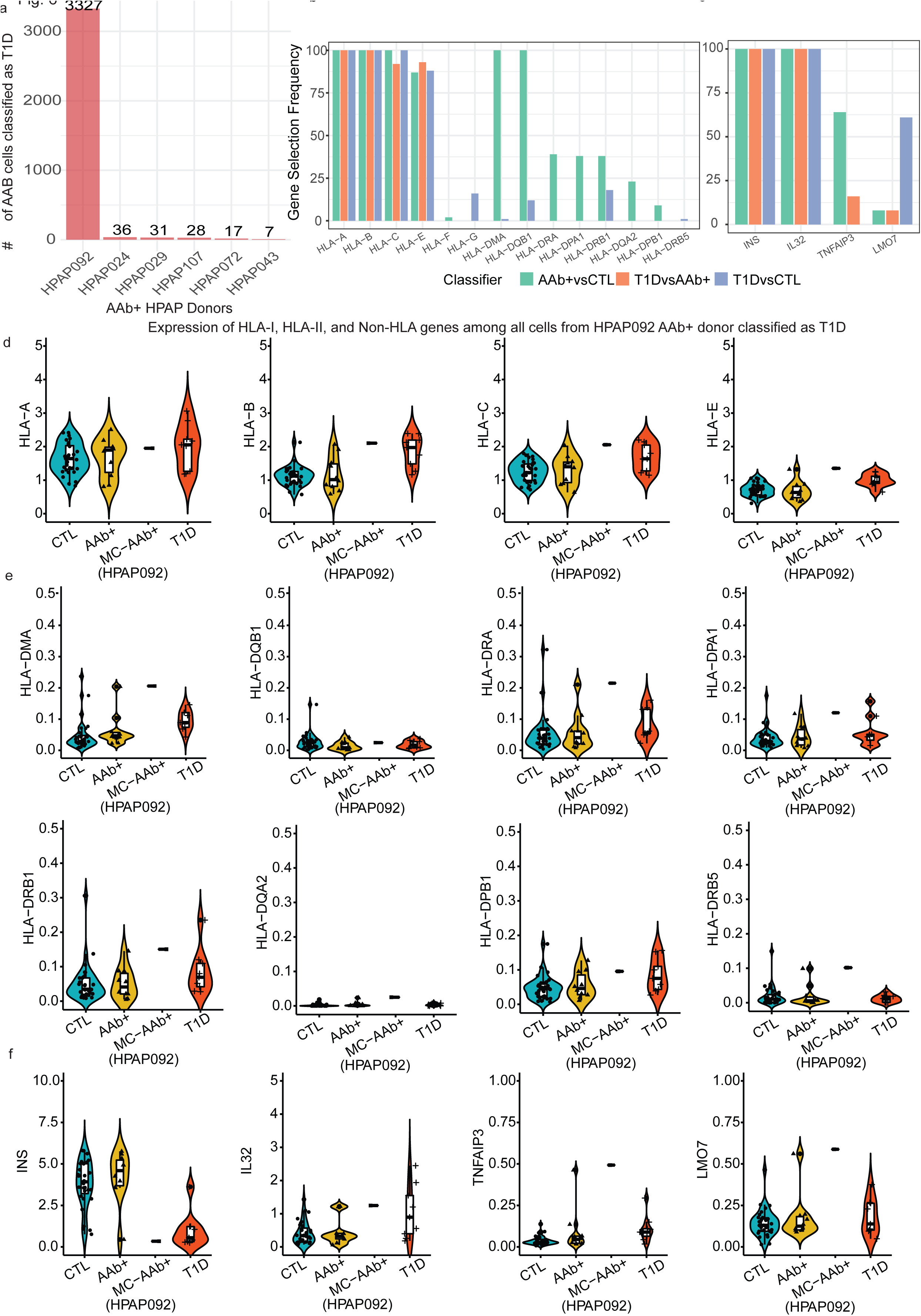
Prediction of AAB+ cells using trained T1D-CTL classifier across all cells together. **a.** Distribution of AAB+ cells predicted as T1D using trained T1D-CTL classifier for all cells **b.** Selection frequency of genes from HLA-I and HLA-II class. **c.** Selection frequency of genes from non-HLA class relevant to T1D. **d.** Comparing the average expression of HLA-I genes among all cells from HPAP092 AAB+ donor classified as T1D. **e.** Comparing the average expression of HLA-II genes among all cells from HPAP092 AAB+ donor classified as T1D. **f.** Comparing the average expression of non-HLA genes among all cells from HPAP092 AAB+ donor classified as T1D.

### Donor-wise classification using the leave-one-out cross-validation strategy

To evaluate the performance accuracy per donor, we applied a unique splitting criterion for testing and training purposes and implemented the leave-one-out cross-validation (LOOCV) strategy. We trained the model based on cells from all donors except one and tested the model’s performance on the one donor left out of the training step. This process was repeated across all donors. Remarkably, in this analysis, the AAb+ donor HPAP092 was classified as T1D across all the annotated cell type comparisons (Fig S4). These results from the LOOCV strategy confirmed of our previous observations (Fig 5a). The mean classification accuracies for each annotated cell type using LOOCV strategy were between 80-90% for most cell types except 72% for the beta cell classifier (Fig S3). This is likely caused by the very few beta cells remaining in several of the donors; for instance, the islets recovered from donors HPAP021 and HPAP022 had only two beta cells each among the total 4,410 and 864 cells analyzed, respectively. This low beta cell count in a subset of T1D donors led to lower overall accuracy by the annotated beta cell classifier of T1D vs. AAb+ using LOOCV. In contrast, when we had employed the annotated beta cell classifier for T1D vs. AAb+ without the LOOCV strategy, all the beta cells were pooled together, leading to high prediction accuracy.

## Discussion

Although substantial progress has been made over the past decades in our understanding of the alterations in the pancreas of T1D patients, the underlying molecular processes of disease progression from healthy to autoantibody positivity to T1D remain to be elucidated fully. Using scRNA-seq islet data from 50 human organ donors, we performed a comprehensive analysis by applying a machine learning approach on the large islet gene expression data from T1D, AAb+, and CTL individuals to understand the disease progression at the single-cell level.

In this study, we focused on the XGBoost method to achieve novel insights into the pre-diabetic and diabetic disease states of pancreatic islets. Sparse read counts are a main characteristic of scRNA-seq data, which is important to consider when performing differential gene expression analysis. The superior performance and robustness of the XGBoost method on high-throughput gene expression studies^28–31^ led to its increased popularity in single-cell biology^32–36^, and this approach remains more powerful compared to other machine learning approaches including neural networks^36^. To the best of our knowledge, there is no literature using the machine-learning approach to identify gene signatures and classification of cells into relevant disease states on single-cell data from T1D donors. We demonstrate excellent performance of XGBoost classifiers built for T1D vs CTL, T1D vs AAb+, and AAb+ vs CTL comparisons across all cells and individual cell types. The average accuracy, sensitivity, and specificity obtained from 100 repetitions were found to be higher in the T1D vs CTL comparison compared to the AAb+ versus CTL, likely because of the high similarity of the latter two states. In addition, the top-selected gene lists obtained for each of these comparisons were found to be upregulated in several key pathways, such as the type 1 diabetes and antigen processing and presentation pathways, with special enrichment seen in beta cells. The genes involved in these pathways belonged to HLA class I; therefore, we measured their individual expression in beta cells and found that they were more highly expressed in T1D donors than in CTL donors.

Increased expression of HLA class I genes in T1D has been reported in the past. Benkahla et al.^20^ reported hyper-expression of HLA class I genes in T1D donors through immunofluorescence staining and microscopic image analysis. Hamilton-Williams et al.^37^ used the non-obese diabetic (NOD) mice model and showed that hyperglycemia was observed in those mice that exhibited higher MHC-I expression in beta cells. Richardson et al.^38^ performed enteroviral capsid protein vp1 staining on a large cohort of neonatal, pediatric control, and T1D groups and observed hyperexpression only in T1D donors. Nejentsev et al.^39^ reported the contribution of *HLA-A*, *HLA-B*, and *HLA-C* towards T1D. In contrast to these studies, Skog et al.^40^ performed staining through immunohistochemical staining (IHC) to measure protein expression and RNA sequencing to measure mRNA expression in non-diabetic controls and T1D patients; however, they reported no changes in the HLA class I genes across these groups. Using imaging mass cytometry, Wang et al found HLA-A, B, and C expression to be upregulated in islets from short but not long duration T1D, and also overexpressed in beta cells in very recent onset disease^41^. Detecting this gene signature in T1D-like AAb+ cells in our study further demonstrate the upregulation of this pathway at early stages of autoimmunity.

A surprising discovery from this study is the observation that a subset of nondiabetic donors positive for islet autoantibodies contain significant numbers of pancreatic cells with gene expression profiles that do not resemble autoantibody negative, nondiabetic controls as expected, but rather those present in the pancreas of T1D individuals. It is tempting to speculate that those AAb+ individuals with a large proportion of ‘T1D-like’ pancreatic cells would have been the ones to progress to the diabetic state more rapidly than those where these cells are not present. Unfortunately, it is impossible to test this hypothesis due to the cross-sectional nature of our study, and the fact that the human pancreas cannot be biopsied safely. Nevertheless, this finding provides strong evidence that the transcriptomic changes that occur during the pathogenesis of T1D are not simply a consequence of the hyperglycemic state; rather, they appear to be an integral part of disease progression.

### Methods

We present an overview of the ML-based XGBoost approach for the classification of pancreatic scRNA-seq islet data from different conditions (Fig. 1a). The complete process included the procurement of human islet tissues, the preparation of a single-cell suspension and 10x Genomics sample processing.

We used R programming language^42^ to perform all the computations, including data pre-processing, ML model building, and downstream calculations and visualizations. The caret^43^, dplyr^44^, and matrix^45^ packages were used for data wrangling tasks and Seurat^46^ for working with the scRNA-seq object. The plots were generated using ggplot2^47^, ggpubr^48^, and cowplot^49^. The ML model was built using XGBoost^9^ for classification purposes, and the gene selection was performed using Ckmeans.1d.dp^50, 51^. All ML computations, including hyperparameter optimization tasks, were performed through parallel computing using #cores between 30 to 100^42^.

### scRNA-seq data description and analysis

The single-cell sequencing (scRNA-seq) experiments were performed using the Single-cell 3’ Reagent v2 and v3 kits from 10X Genomics. The libraries were processed using 10X Genomics Cell Ranger v6.1 software which aligns the reads and generates feature-barcode matrices (3’ gene expression data) using a collection of several pipelines^52^. The experimental details and pre-processing steps for all the samples from pancreatic-islet data were followed as described previously^11, 53^. We first obtained HPAP samples from different biological conditions such as auto-antibody positive (AAB+; n=10; cells=36,244), type-1 diabetes (T1D; n=9; cells=34,524), and healthy controls (CTL; n=31; cells=123,102). The AAB+ samples were determined based on the AAB+ screening test to measure the levels of antibodies against glutamic acid decarboxylase (GAD) to determine positivity^11, 53^. The combined feature-barcode matrix, including cells from all HPAP samples, added to a total of 193,870 cells. We pre-processed the raw data following the protocol described previously^11, 53^, excluding type-2 diabetes (T2D) samples. We also performed an additional quality control step by removing the mitochondrial and ribosomal reads from the raw data. We then used the exact pipeline described in^11, 53^ for downstream analysis using Seurat v4.1.0^54^ for creating the single-cell object, scDblFinder v1.8.0^55^ for removing doublets from the data, SingleCellExperiment v1.16.0^56^ for data wrangling, sctransform v0.3.3^57^ for data transformation through normalization and scaling, and finally scSorter v0.0.2^12^ for cell-type annotations. The final processed and annotated scRNA-seq data object contained a total of 169,027 cells and 30,002 genes for which transcripts could be detected in at least on cell.

### Machine learning classification network architecture and training protocol

The annotated scRNA-seq data includes 50 HPAP donor samples with 169,027 cells and 30,002 genes from three groups of CTL (n=31), AAB+ (n=10), and T1D (n=9). The XGBoost machine learning framework is constituted of inner and outer loops. In the outer loop, we first randomly split the pre-processed single-cell data into training and testing sets with 70% and 30% of HPAP samples, respectively. In the inner loop, the training was subjected to hyperparameter tuning, and a 5-fold cross-validation was performed across 200 sub-training model iterations to select the best model with a minimum error rate. We repeated this hyperparameter optimization procedure ten times to select the best weights and parameters. The final best weights and parameters obtained were applied to the complete 70% of the training data matrix to obtain the final best training model. Lastly, we applied 30% of the test data matrix on the best-trained model to evaluate predictions. We used the evaluation metrics of accuracy, sensitivity, and specificity on the test data to measure the performance of the model. Along with these metrics, we also obtained the ranked list of genes, also called the best features, from the best-trained model.

To increase the robustness of our approach and attain reliable results, we repeated this entire process which consists of randomly splitting the single-cell data into training (70%) and testing (30%) by using random sampling without replacement in the outer loop for 100 iterations. The mean and SD classification accuracy, sensitivity, and specificity across 100 XGBoost iterations were calculated for the final average performance evaluation. For the final significant gene selection, we built a matrix of genes X iterations. The ranked list of genes obtained in each outer loop iteration was added to the matrix. We counted the gene occurrences across all 100 iterations and ranked them accordingly. The final significant gene list consisted of the list of genes with the count showing how many times they were selected in the best-trained model in 100 iterations. The complete workflow of the XGBoost scRNA-seq ML framework is shown in Fig. 2a.

### XGBoost method

eXtreme Gradient Boosting (XGBoost)^9^ is an extension of gradient-boosted decision trees (GBDT) and is specifically optimized to provide faster computations through parallel and distributed computing. This ensemble learning method uses a boosting approach to improve prediction accuracy by building many aggregated trees to form a single consensus prediction model^58^. The least squares loss function is used to reduce the loss^59^. The XGBoost method creates trees, and the residuals obtained from previous trees are given as input to the subsequent tree, which improves the overall prediction by modeling the errors. For each tree sequence, a subsample of training data is randomly drawn without replacement from the entire training data and is used to fit the tree and compute the model update. Additional trees are not allowed to be added after reaching the pre-specified maximum threshold or if the models converges and the performance does not improve which helps to avoid over parametrization. Overall, this highly effective scalable tree boosting system was originally proposed for sparse data and weighted quantile sketch for approximate tree learning^9^. XGBoost method is applied in a wide range of applications such as regression, classification, ranking, and user-defined prediction problems.

### Feature importance score

The trained XGBoost model automatically provides a feature importance score^60^. The score indicates how important the feature (gene in this study) was used for the model’s prediction. The feature importance score was obtained for each trained model. We selected the genes with non-zero importance scores for all models. We obtained the ranked lists of genes across 100 repetitions of train-test splits.

### Hyperparameter optimization (HPO)

The XGBoost method uses several parameters to control the bias-variance tradeoffs. The tree-based boosting models could suffer from overfitting; however, the XGBoost method provides several parameters, such as maximum tree depth, minimum leaf weight, and minimum split gain, which helps to avoid overfitting. Additionally, it adds randomness to the model during the training phase, making it more robust to noise. Following are the list of parameters set during cross validation on training, *max_depth-* the maximum depth of the tree between 3 to 7, *gamma-* represents the minimum loss reduction required to make a further partition on a leaf node of the tree and it was set between 0 to 0.2, *eta-* the step size of each boosting step was set using random uniform distribution between 0.01 and 0.3, the *min_child_weight-* was set to 30 as large number is usually conservative, *subsample-* the subsample ratio will randomly sample the training data prior to growing trees which will avoid overfitting was set between 0.5 to 0.8 using random uniform distribution, *colsample_bytree*-was also set between 0.5 to 0.8, *cv.nrounds*-200 represents the maximum number of rounds for cross validation, and the *cv.nfold*=5 shows the five-fold cross validation, *early_stopping_rounds*=100 which helps the model to stop training further if the best performance was achieved, *eval_metric=*logloss was used to minimize the loss, *Binary=*logistic was used for performing logistic regression for binary classification and output the probabilities, nthreads=30, for parallelization of all the tasks. In each inner loop, we iterated ten times over the list of above-described parameters to identify the best parameters and used them to train the XGBoost model and test the predictions through the test dataset. Lastly, this procedure was repeated 100 times in the outer loop, and the average performance metrics were calculated.

### Leave one out cross-validation strategy (LOOCV)

We used a special case of k-fold cross-validation called LOOCV where for each sample in the dataset, removing that sample (testing), training was performed on all the remaining samples. We used the LOOCV instead of previous criteria of training (70%) and testing (30%), where random sampling without replacement was considered in the outer loop for 100 iterations. The same parameters previously mentioned in HPO pipeline were used here for optimizing the model during training. The LOOCV helps to classify disease vs. normal group at subject level where the subject being tested is classified as either disease or normal.

### Evaluating performance

We measure the performance of XGBoost models using averages of accuracy, sensitivity, and specificity obtained across 100 iterations. The equations for these metrics with respect to true positive (TP), true negative (TN), false negative (FN), and false positive (FP) are as follows:

The classification accuracy of the model is defined as the ratio of correctly predicted instances (TP + TN) to the total number of instances in the dataset (TP +TN + FP + FN)

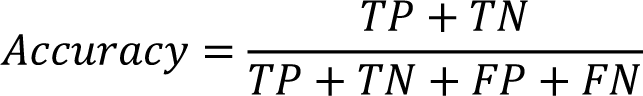

Sensitivity measures the model’s ability to correctly identify positive instances (TP) out of all the instances that are actually positive (TP + FN).

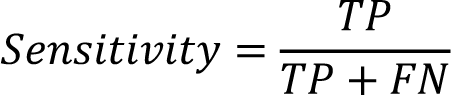

And finally, Specificity shows the model’s ability to correctly identify negative instances (TN) out of all the instances that are actually negative (TN + FP).

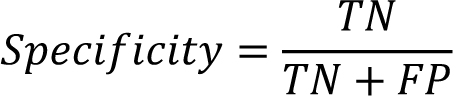

### Gene selection and pathway enrichment analysis

We obtained the ranked lists across 100 repetitions of train-test splits (Methods, section-Feature Importance score) and followed two strategies with different criteria to obtain list of ranked gene lists:

1. **Ranked gene selection:** We aggregated the ranked lists across 100 repetitions by preserving the ranks by applying the robust rank aggregation (RRA)^14^ method to obtain the final ranked list of genes in a given comparison. RRA is a statistical technique used to combine rankings from multiple sources into a single, aggregated ranking that is robust to noise, inconsistencies, and outliers in the individual rankings. The RRA method assigns scores in terms of P-values for each gene to determine the significance level. We used genes with P-value < 0.05 for the final ranked list of genes for all comparisons (Supplementary Tables S2, S3, and S4).
2. **Unranked gene selection:** We aggregated the ranked lists across 100 repetitions by ignoring the ranks and focusing on the number of times a gene was selected across 100 repetitions. For example, in beta cells of T1D vs. CTL classifier results, if gene *INS* is selected 80 times among 100 repetitions, then a selection frequency score of 80 was assigned to *INS*. The final ordered list of genes with score as gene selection frequency among 100 repetitions for all comparisons (Supplementary Tables S5, S6, and S7).

For the ranked list of genes, we performed pathway analysis using the ClusterProfiler^61^ tool. The clusters were created for the gene ontology (GO) biological pathways (BP) and Kyoto Encyclopedia of Genes and Genomes (KEGG) pathways based on the ranked list of genes across each comparison. Our analysis determined the GO biological processes BP and KEGG pathways with an adjusted P-value < 0.05 as statistically significant (Supplementary Tables S8, S9, and S10). Next, we created a shared list of ranked genes using the RRA method and performed GO and KEGG pathway analysis using clusterProfiler^61^ and Metascape^62^, based on the ranked gene lists obtained for different cell types. The ranked gene lists across different cell types in each comparison (Supplementary Tables S2, S3, and S4) were given as input to the RRA method. The obtained shared list of ranked genes was called “RRA_combined” and reported along with the corresponding GO and KEGG pathways (Supplementary S14, S15, and S16).

For the unranked list of genes, the publicly available databases such as the ‘database for annotation, visualization, and integrated discovery’ (DAVID, version 6.8) bioinformatics web server^63^ was used to identify various biological pathways of significant genes through a set of functional annotation tools. We first filtered high confidence genes by selection genes with a selection frequency of greater than 50 and used the GO^64, 65^ and the KEGG^66–68^ databases for pathway enrichment analysis in the DAVID database. The GO classifies the gene functionalities into three categories: biological processes (BP), cellular component (CC), and molecular function (MF), and the KEGG database provides an overview of high-level gene functions and biological signaling pathways. Our analysis determined the GO and KEGG terms with false discovery rate (FDR) < 0.05 as statistically significant (Supplementary S11, S12, and S13).

### Protein-protein interaction networks and gene modules selection

The Search Tool for Retrieval of Interacting proteins database (STRINGdb v11)^69^ was used to identify the various protein-protein interaction networks (PPI) of significant gene lists based on the medium confidence score of 0.7. The loaded PPI network from STRINGdb^69^ was analyzed using the open-source Cytoscape^70, 71^ tool. Cytoscape is a bioinformatics software used for visualizing and integrating highly complex PPI networks through several supported plugins. The StringApp^70^ within Cytoscape is used to load the raw PPI network, and the network analyzer^72^ plugin is used to measure the degree of interaction between nodes and display the up/down-regulated genes. Finally, we applied the molecular complex detection (MCODE)^73^ clustering-based algorithm for further splitting the network into modules/clusters that helped identify the densely connected regions. We used the default parameters (degree cutoff=2, node score cut-off=0.2, k-core=2, and max.depth=100) to filter and identify key clusters in the network. We selected the top modules from each analysis group to show the degree of interactions.

## Materials availability

This study did not generate new unique reagents.

## Data availability

The pancreatic islet sequencing data can be found in PANCDB (https://hpap.pmacs.upenn.edu/analysis).

## Author contributions

Data generation, software, A.R.P.; resources, A.R.P.; writing—original draft preparation, A.R.P.; writing— review and editing, A.R.P. G.V.; K.H.K and R.B.F review and editing. All authors have read and agreed to the published version of the manuscript.

## Competing interests

The authors declare no competing interests.

## Additional information

**Correspondence** and requests for materials should be addressed to Lead Contact, Golnaz Vahedi.

## Supplementary Figures Legends

**Supplementary Figure S1.** Significant KEGG pathways (FDR < 0.05) based on ranked genes obtained from T1D vs. AAb+ comparison across Unannotated all cells and annotated different cell types (Table S3).

**Supplementary Figure S2.** Significant KEGG pathways (FDR < 0.05) based on ranked genes obtained from AAb+ vs. CTL comparison across Unannotated all cells and annotated different cell types (Table S4).

**Supplementary Figure S3.** Stacked bar chart showing the donor wise classification accuracy based on LOOCV approach for T1D vs. AAb+ classifier across annotated cell types.

**Supplementary Figure S4.** Stacked bar chart showing the donor wise percentage of AAb+ cells classified as T1D or AAb+ across annotated cell types.

## Table Legends

**Table S1.** Word document showing the classification performance for different pairwise comparisons (T1D vs. CTL, T1D vs. AAb+, and AAb+ vs. CTL) across unannotated all cells and annotated cell types.

**Table S2.** Ranked significant gene lists obtained through RRA approach (P-value < 0.05) in unannotated all cells and annotated cells (Acinar, Alpha, Beta, Delta, Ductal, Immune, Endothelial, and Stellates) for T1D vs. CTL pairwise comparison.

**Table S3.** Ranked significant gene obtained through RRA approach (P-value < 0.05) in unannotated all cells and annotated cells (Acinar, Alpha, Beta, Delta, Ductal, Immune, Endothelial, and Stellates) for T1D vs. AAb+ pairwise comparison.

**Table S4.** Ranked significant gene lists obtained through RRA approach (P-value < 0.05) in unannotated all cells and annotated cells (Acinar, Alpha, Beta, Delta, Ductal, Immune, Endothelial, and Stellates) for AAb+ vs. CTL pairwise comparison.

**Table S5.** Unranked significant gene lists with gene selection frequency in unannotated all cells and annotated cells (Acinar, Alpha, Beta, Delta, Ductal, Immune, Endothelial, and Stellates) for T1D vs. CTL pairwise comparison.

**Table S6.** Unranked significant gene lists with selection frequency in unannotated all cells and annotated cells (Acinar, Alpha, Beta, Delta, Ductal, Immune, Endothelial, and Stellates) for T1D vs. AAb+ pairwise comparison.

**Table S7.** Unranked significant gene lists with selection frequency in unannotated all cells and annotated cells (Acinar, Alpha, Beta, Delta, Ductal, Immune, Endothelial, and Stellates) for AAb+ vs. CTL pairwise comparison.

**Table S8.** GO-BP and KEGG pathways for unannotated all cells and annotated cells (Acinar, Alpha, Beta, Delta, Ductal, Immune, Endothelial, and Stellates) using clusterProfiler in T1D vs. CTL pairwise comparison based on ranked gene lists based on RRA (P-value < 0.05) from Table S2.

**Table S9.** GO-BP and KEGG pathways for unannotated all cells and annotated cells (Acinar, Alpha, Beta, Delta, Ductal, Immune, Endothelial, and Stellates) using clusterProfiler in T1D vs. AAb+ pairwise comparison based on ranked gene lists based on RRA (P-value < 0.05) from Table S3.

**Table S10.** GO-BP and KEGG pathways for unannotated all cells and annotated cells (Acinar, Alpha, Beta, Delta, Ductal, Immune, Endothelial, and Stellates) using clusterProfiler in AAb+ vs. CTL pairwise comparison based on ranked gene lists based on RRA (P-value < 0.05) from Table S4.

**Table S11.** GO (GO-BP, GO-MF, GO-CC) and KEGG pathways for unannotated all cells and annotated cells (Acinar, Alpha, Beta, Delta, Ductal, Immune, Endothelial, and Stellates) using DAVID in T1D vs. CTL pairwise comparison based on unranked gene lists based on gene selection frequency > 50 from Table S5.

**Table S12.** GO (GO-BP, GO-MF, GO-CC) and KEGG pathways for unannotated all cells and annotated cells (Acinar, Alpha, Beta, Delta, Ductal, Immune, Endothelial, and Stellates) using DAVID in T1D vs. AAb+ pairwise comparison based on unranked gene lists based on gene selection frequency > 50 from Table S6.

**Table S13.** GO (GO-BP, GO-MF, GO-CC) and KEGG pathways for unannotated all cells and annotated cells (Acinar, Alpha, Beta, Delta, Ductal, Immune, Endothelial, and Stellates) using DAVID in AAb+ vs. CTL pairwise comparison based on unranked gene lists based on gene selection frequency > 50 from Table S7.

**Table S14.** Shared list of ranked genes called RRA_combined obtained by combining gene lists from Table S2 using RRA approach in T1D vs. CTL. The corresponding GO and KEGG pathways for shared ranked genes are also shown.

**Table S15.** Shared list of ranked genes called RRA_combined obtained by combining gene lists from Table S3 using RRA approach in T1D vs. AAb+ The corresponding GO and KEGG pathways for shared ranked genes are also shown.

**Table S16.** Shared list of ranked genes called RRA_combined obtained by combining gene lists from Table S4 using RRA approach in AAb+ vs. CTL The corresponding GO and KEGG pathways for shared ranked genes are also shown.

## Notes

### Competing Interest Statement

The authors have declared no competing interest.

